# Identification of quercetin from fruits to immediately fight zika

**DOI:** 10.1101/074559

**Authors:** Amrita Roy, Liangzhong Lim, Jianxing Song

## Abstract

Zika virus is spread mainly by the bite of an infected mosquito, which can be passed from a pregnant woman to her fetus, thus leading to birth defects including more than microcephaly. It has been recently estimated that one-third of the world population will be infected by Zika in the near future, but unfortunately so far there is no vaccine or medicine for Zika. In particular, the special concern on the vaccine treatment to Zika and Dengue arising from antibody-dependent enhancement strongly emphasizes the key role of its NS2B-NS3 protease (NS2B-NS3pro) as a target for anti-Zika drug discovery/design due to its absolutely-essential role in viral replication.

In response to the current global health emergency triggered by the Zika outbreak, we successfully obtained several active forms of Zika NS2B-NS3pro and further attempted to discover its inhibitors from eatable plants and traditional herbal medicines to immediately fight Zika. Here, for the first time, we discovered that quercetin, a flavonoid extensively existing in many fruits and vegetables, effectively inhibits Zika NS2B-NS3pro. We further quantify its inhibitory activity with IC_50_ of 26.0 ± 0.1 µM; and *K*_i_ of 23.0 ± 1.3 µM. As quercetin has been extensively found in fruits, vegetables, leaves and grains, our discovery would benefit the public to immediately fight Zika.

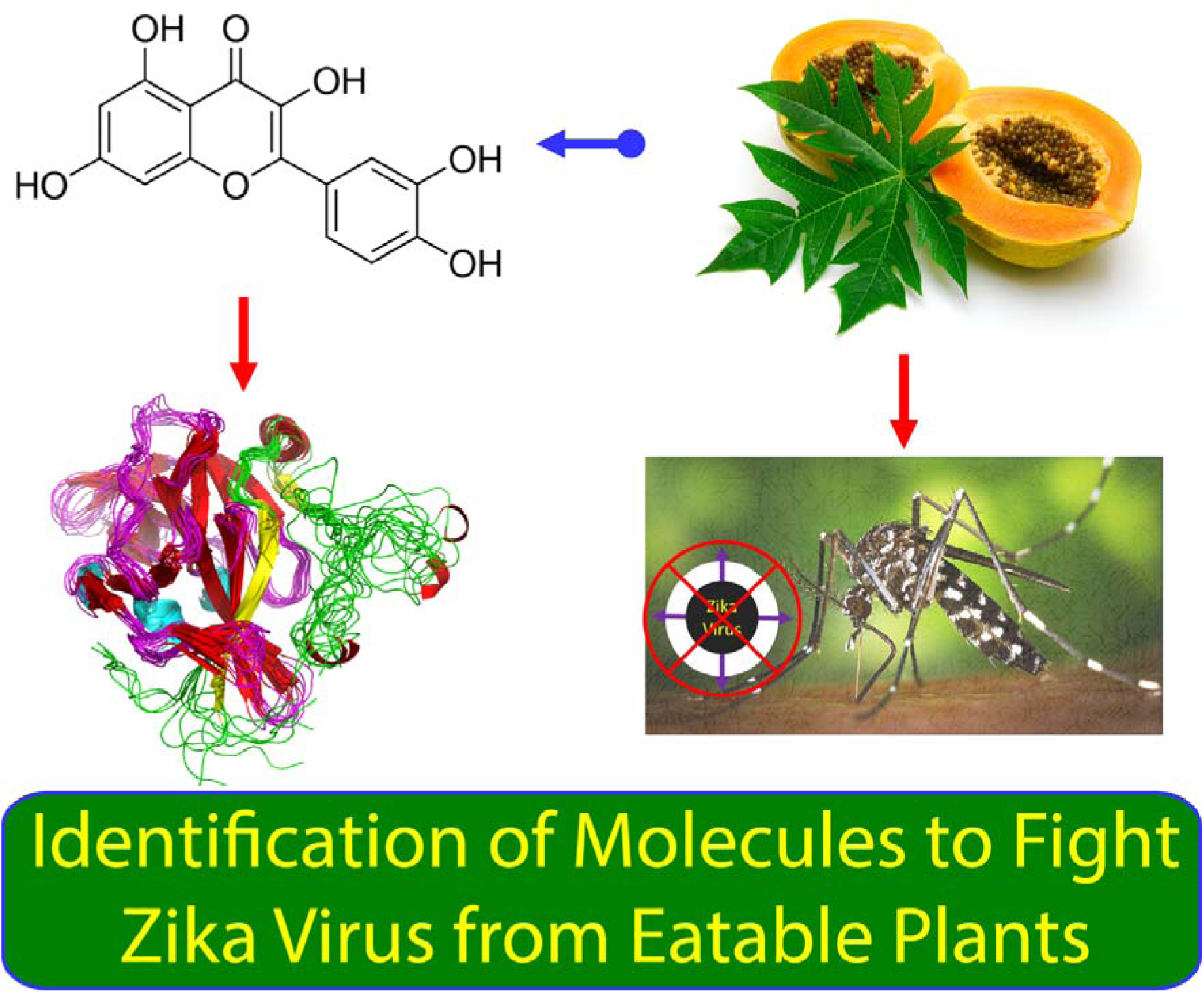

## Introduction

Zika virus was originally isolated from a sentinel rhesus monkey in the Zika Forest of Uganda in 1947 (1), which is transmitted to humans by Aedes species mosquitoes. Since 2007, large epidemics of Asian genotype Zika virus have been reported around the world (2–4). Recently it has been estimated that one-third of the world population will be infected in the near future (5). Most seriously, Zika infection has been found to be associated with serious sequelae such as Guillain-Barré syndrome, and microcephaly in newborn infants of mothers infected with Zika virus during pregnancy (6–9), and consequently WHO has declared a public health emergency for Zika virus (10). Zika virus represents a significant challenge to the public health of the whole world but unfortunately there is no effective vaccine or other therapy available so far.

Zika virus with a single stranded, positive sense RNA genome of 10.7 kb belongs to the flavivirus genus, which also contains Dengue, yellow fever, West Nile, Japanese encephalitis, and tick-borne encephalitis viruses (4,11). Zika virus shares a high degree of sequence and structural homology with other flaviviruses particularly Dengue virus, thus resulting in immunological cross-reactivity (7). As such, Zika was proposed as the fifth member of the Dengue serocomplex. Seriously, the current Zika outbreaks are largely localized within dengue-endemic areas, it is thus possible that preexisting dengue-induced antibodies may enhance Zika infection by antibody-dependent enhancement (ADE), a factor that makes the vaccine approaches extremely challenging (7).

Zika genome is translated into a single ~3,500-residue polyprotein, which is further cleaved into 3 structural proteins and 7 non-structural proteins (11). The correct processing of the polyprotein is essential for replication of all flaviviruses, which requires both host proteases and a viral NS2B-NS3 protease (NS2B-NS3pro) (11–18). As a consequence, the flaviviral NS2B-NS3pro has been well-established to be key targets for developing antiviral drugs (11–18). In particular, the unique concern on the vaccine treatment to Zika and Dengue strongly emphasizes the irreplaceable role of the Zika protease as a target for antiviral drug discovery/design.

As facilitated by our previous studies on Dengue NS2B-NS3pro (12), we started to work on Zika NS2B-NS3pro immediately after the outbreak of Zika and now successfully obtained several active forms of recombinant Zika NS2B-NS3pro. Furthermore, we attempted to discover its inhibitors from eatable plants and traditional herbal medicines to fight Zika with no delay. Here we release the result that quercetin, a flavonoid existing in many fruits and vegetables, inhibits Zika NS2B-NS3pro, and subsequently determined its inhibitory activity. For the first time, our study reveals that many fruits, vegetables, leaves and grains, as well as herbals containing quercetin are expected to have anti-Zika activity, and therefor can join in to combat Zika immediately.

## Results and discussion

### Cloning, expression and purification of the active Zika NS2B-NS3pro

Based on the sequence alignment with NS2B and NS3pro of the Dengue serotype 2 we previously studied (12), the corresponding Zika sequences were identified for the NS2B and NS3pro of the Asian Zika virus. From synthetic genes with *E. coli* preferred codons, we amplified and subsequently cloned the DNA fragments into His-tagged expression vectors. The recombinant proteins were expressed in BL 21 cells and then purified by Ni^2+^-affinity chromatography, followed by further purification with FPLC gel-filtration chromatography. Recombinant protease samples were checked by SDS-PAGE, molecular weights verification with ESI-MS and protein sequencing with time-of-flight-mass spectrometer (Applied Biosystems). Protein concentration was determined by the UV spectroscopic method with 8 M urea (19).

All enzymatic/inhibitory experiments were performed in 50 mM Tris-HCl at pH 8.5, with a Cary Eclipse fluorescence spectrophotometer in triplicate and data are presented as mean ± SD. The Zika NS2B-NS3pro we obtained is fully active as judged from its enzymatic activity on a fluorescent substrate Bz-nKRR-AMC (12).

### Identification of quercetin as an inhibitor

Quercetin purchased from Sigma-Aldrich (Fig 1A) is a flavonoid extensively existing in many fruits, vegetables, leaves and grains such as *Carica papaya* (Fig 1B). When Zika NS2B-NS3pro at 50 nM (Fig 1C) was pre-incubated with quercetin at different concentrations, the enzymatic activity was inhibited at a dose-dependent manner (Fig 2A), indicating that quercetin is an inhibitor of Zika NS2B-NS3pro.

**Fig 1.**
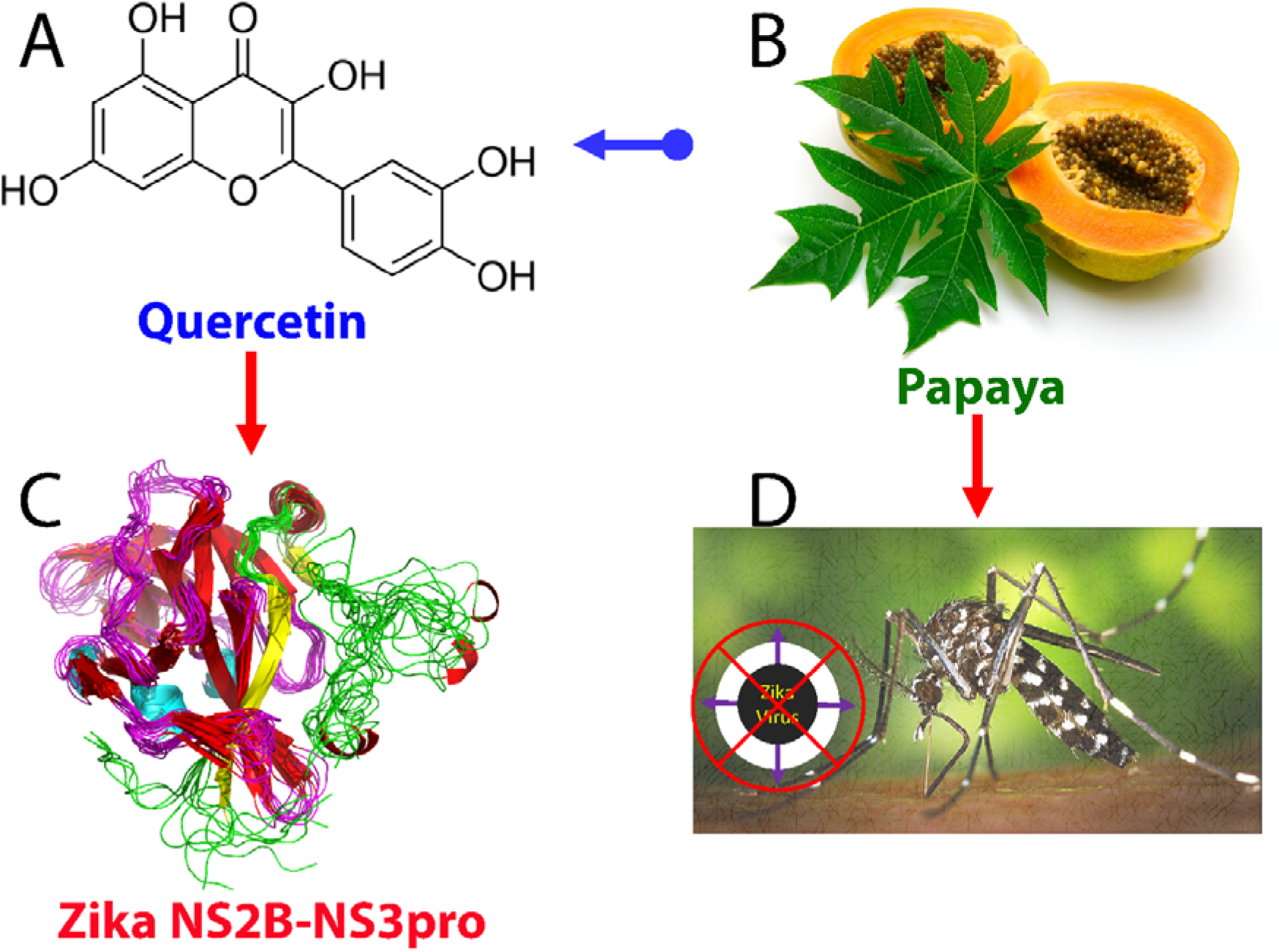
Identification of an inhibitor of Zika NS2B-NS3pro from eatable plants. (A). Chemical structure of quercetin extensively existing in many fruits and vegetables including Papaya (B). Here we identified quercetin to be an inhibitor of Zika NS2B-NS3pro (C); thus suggesting that fruits and vegetables containing quercetin have anti-Zika activity (D).

**Fig 2.**
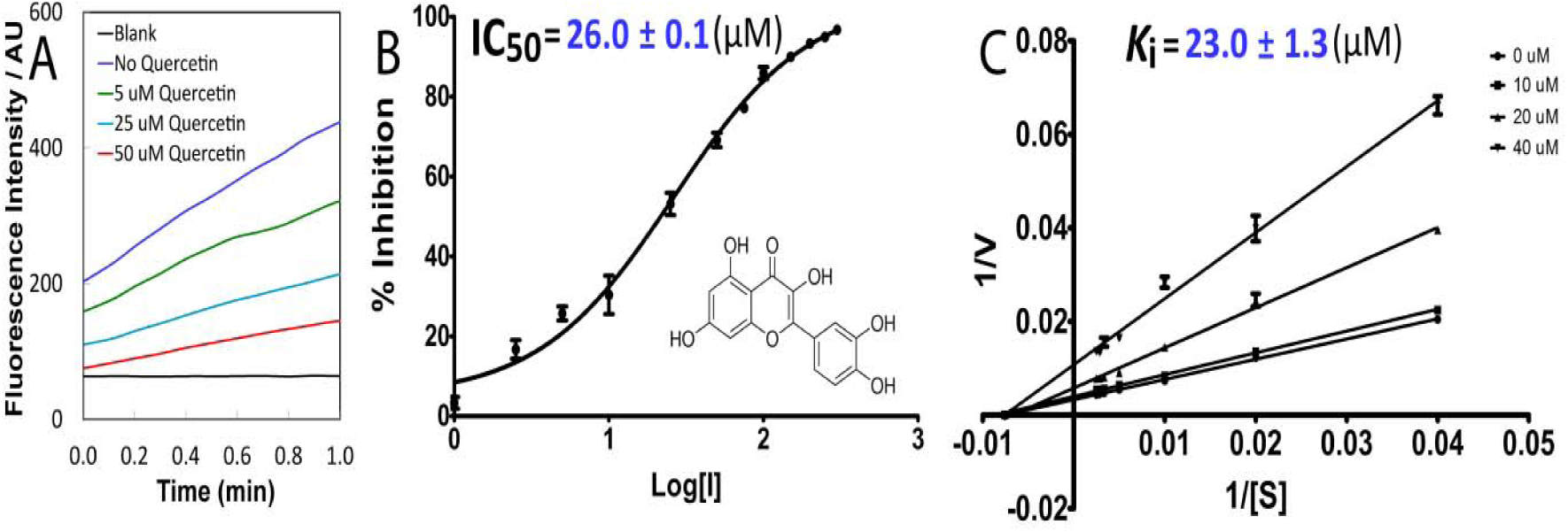
Characterization of inhibitory effects of quercetin. (A) The tracings of the fluorescence intensity within 1 min for Zika NS2B-NS3pro in the presence of quercetin at different concentrations. (B) Fitting curve for IC_50_ of quercetin. (C) The Lineweaver-Burk plot for determing *K* of quercetin. [S] is the substrate concentration; V is the initial reaction rate. Both curves were generated by the program GraphPad Prism 7.0.

Therefore we further determined the IC_50_ and *K*_i_ values of quercetin. Briefly, for determination of IC_50_, the protease at 50 nM was pre-incubated at 37 °C for 30 min with quercetin at various concentrations and the reaction was initiated by adding Bz-nKRR-AMC to 250 μM. The obtained data (Fig 2B) were fitted to be 26.0 ± 0.1 µM by GraphPad Prism 7.0 (20)

For *K*i determination, the assay was performed with different final concentrations of both quercetin and substrate. The protease at 50 nM was pre-incubated with quercetin at different concentrations for 30 min at 37 ºC. Subsequently, the reaction was initiated by addition of the corresponding concentration series of the substrate. The *K*i was obtained to be 23.0 ± 1.3 µM by fitting the data (Fig 2C) in the non-competitive inhibition mode (20). It is interesting to find that quercetin acts as a non-competitive inhibitor for Zika NS2B-NS3pro. This clearly suggests that quercetin is an allosteric inhibitor which binds to the site having no direct overlap with the substrate binding pocket of Zika NS2B-NS3pro. We are currently focused on decoding whether quercetin in fact acts as a dynamically-driven allosteric inhibitor as previously observed on other viral proteins (21–25). Furthermore, we also attempted to utilize NMR spectroscopy to facilitate the optimization of the inhibitors (26–31).

Quercetin is a flavonoid extensively found in many fruits, vegetables, leaves and grains, which have been used as an ingredient in supplements, beverages, or foods. It has been previously documented that quercetin has beneficial health effects ranging from antioxidant to nutraceutical (32). Recently it has also been shown that herbal medicines containing quercetin or quercetin-like molecules showed significant antiviral activities on SARS and Dengue viruses (33–35). Here, for the first time, we provide the experimental evidence that quercetin is also an inhibitor of Zika NS2B-NS3pro, which is absolutely essential for cleaving the Zika polyprotein into functional subunits. Therefore, quercetin-containing fruits and vegetables can join in to fight Zika virus right away. Moreover, quercetin can also serve as a starting point for further design of inhibitors of high affinity and specificity for fighting Zika.

## Conclusion

For the first time, our study discloses that quercetin is an inhibitor of Zika NS2B-NS3 protease, which consequently should have capacity in inhibiting Zika replication. In particular, with consideration of the fact that quercetin has been extensively found in many fruits, vegetables, leaves and grains, our discovery would benefit the public to immediately fight Zika.

## Acknowledgement

This study is supported by Ministry of Education of Singapore (MOE) Tier 3 Grant R-154-002-580-112; Tier 2 Grant MOE2015-T2-1-111 and National Research Foundation of Singapore (NRF) R-154-002-529-281 to Jianxing Song. The funders had no role in study design, data collection and analysis, decision to publish, or preparation of the manuscript.

## Author Contributions

Conceived the idea: JXS; Performed the experiments: AR LZL; Prepare figures and wrote the paper: JXS.

